# Developing a low-cost milliliter-scale chemostat array for precise control of cellular growth

**DOI:** 10.1101/223594

**Authors:** David Skelding, Sam Hart, Thejas Vidyasagar, Alexander E. Pozhitkov, Wenying Shou

**Affiliations:** Division of Basic Sciences, Fred Hutchinson Cancer Research Center, Seattle, WA; University of Washington, Seattle, WA

## Abstract

Multiplexed milliliter-scale chemostats are useful for measuring cell physiology under various degrees of nutrient limitation and for experimental evolution. In each chemostat, fresh medium containing a growth rate-limiting metabolite is pumped into the culturing chamber at a constant rate, while culture effluent exits at an equal rate. Although such devices have been developed by various labs, key parameters - the accuracy and precision of flow rate and the operational range - are not explicitly characterized. Here we report the development of multiplexed milliliter-scale chemostats where flow rates for eight chambers can be independently controlled to vary within a wide range, corresponding to population doubling times of 3~ 13 hours. Importantly, flow rates are precise and accurate without the use of expensive feedback systems. Among the eight chambers, the maximal coefficient of variation in flow rate is less than 3%, and average flow rates are only slightly below targets, *i.e.*, 3-6% for 13-hour and 0.6-1.0% for 3-hour doubling times. This deficit is largely due to evaporation and should be correctable. We experimentally demonstrate that our device allows accurate and precise quantification of population phenotypes.

## Introduction

Continuous-culturing devices are useful for measuring microbial phenotypes in a constant environment or for performing experimental evolution. Unlike batch culturing where an exponentially-growing population eventually enters stationary phase due to nutrient exhaustion or waste accumulation, continuous-culturing devices constantly supply nutrients and remove waste products.

Commonly used continuous-culturing devices include turbidostats and chemostats. A turbidostat maintains a population at a constant turbidity in a nutrient-rich environment ^1^. For example, in Klavins lab’s multiplexed turbidostats ^2^, culture turbidity in each chamber is measured via a laser beam and a light detector. Once turbidity exceeds a target value, custom software directs the flow of fresh medium using pinch valves and a syringe pump (Fig 1B). The input medium is mixed into the chamber to create a uniform environment, and excess volume is forced out to waste via positive pressure exerted by the inflow of filtered and humidified air. Because current turbidity is always compared to the target turbidity in a feedback loop, turbidity remains steady despite any transient fluctuations in flow rate.

**Fig. 1.**
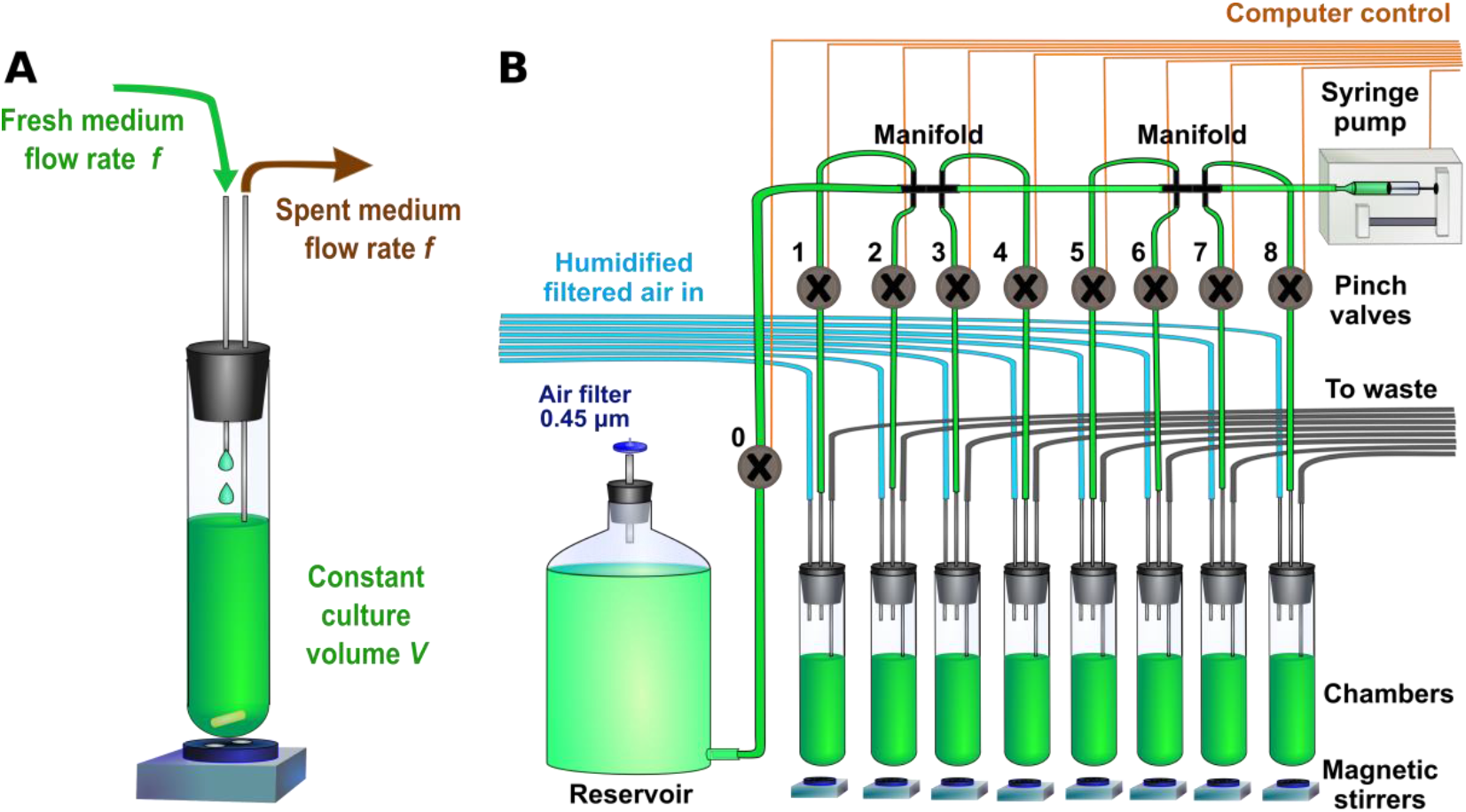
Chemostat and a turbidostat-chemostat array. (**A**) In a chemostat, fresh medium with a growth rate limiting metabolite flows into chamber at a constant rate *f*, and is mixed with the contents of the chamber. Excess culture effluent exits at the same rate, thus maintaining a constant volume *V*. (**B**) In the Klavins lab 8-chamber design, a computer-controlled syringe pump is used for all liquid dispenses. Pinch valves (grey circles) are used to open and close tubing. Excess volume is forced out to waste using positive pressure created by the inflow of humidified and filtered air (blue lines). Mixing is accomplished using magnetic stirrers. In the Klavins lab setup, each cycle proceeds as the following. First, only Valve 0 opens, and sufficient fresh medium is drawn from the reservoir for all eight chambers and for backlash correction (Supplementary Fig 3). Next, the backlash volume is returned to reservoir and Valve 0 is closed. Subsequently, Valve 1 to Valve 8 opens one at a time in sequence, and an appropriate amount of medium is dispensed into each Chamber.

In contrast, a chemostat creates a nutrient-limited environment where the population is forced to grow at a constant, pre-determined rate slower than the maximal growth rate. Specifically, a chemostat chamber (Fig 1A) contains a culture of a fixed volume *V*. A medium containing a limiting metabolite is added at a constant flow rate *f*(ml/h). The culture effluent is removed from the chamber at the same rate *f*, thereby maintaining a constant volume. Mathematically ^3^, live population density *N* satisfies

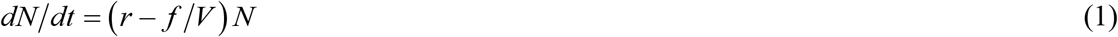

where *r* (h^-1^) is the net growth rate (birth rate minus death rate), *f* is the flow rate, and *f/V* is the dilution rate (h^-1^). At steady state, the growth rate and the dilution rate are equal:

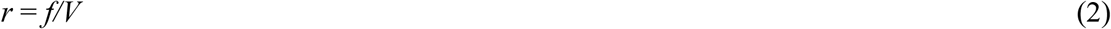

*M*, the concentration of limiting metabolite in culturing vessel satisfies:

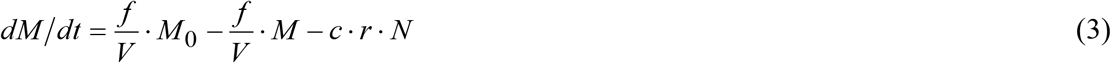

where *M_0_* is the metabolite concentration of inflow fresh medium and *c* is the amount of metabolite consumed per net growth of one cell. Thus, steady-state cell density may be controlled by *M_0_*:

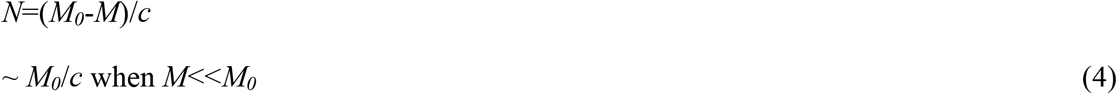

Chemostats are useful for measuring cell starvation phenotypes and for evolving cells under nutrient limitation. Multiple chemostats are often desired to test experimental repeatability or different degrees of nutrient limitation. Commercially-available chemostats are expensive and are usually geared toward large culture volumes (liter-scale instead of a few milliliters). Miniature multi-chamber turbidostat/chemostat devices have been constructed previously ^4–6^. However, the consistency and accuracy of flow rates and the operational ranges in these designs are not explicitly uncharacterized. For example, in Matteau *et al* device ^5^, flow rate is mediated via valve opening time. At the shortest valve opening time they tested (1 sec), actual dispensed volume (~0.49 ml) seemed to deviate from the regression by tens of percent. Moreover to realize 10h doubling time in a 20 ml culturing chamber, the flow rate will be 1.39 ml/hr. This translates to around three 1s-dispenses per hour. Thus, flow is rather discontinuous at slow doubling times.

Precise flow rates may be achieved via a feedback mechanism. For example, one could use the decline rate of medium reservoir weight to measure past flow rate, and adjust future flow rate accordingly. However, this would require each chamber to have its own reservoir and scale, which is not easily scalable. Without any flow rate feedback control, chemostat flow rates were quite variable and could deviate significantly from the target as we tried to run Klavins lab’s turbidostats as chemostats. Here, we have modified Klavins lab multiplexed turbidostats to offer the additional functionality of multiplexed chemostats with consistent, accurate, and independently controllable flow rates.

## Results and Discussions

### Variable dilution rates in multiplexed turbidostats

We tested the original Klavins design by running all eight chambers at the same target flow rate. We measured flow rate in each chamber by quantifying media accumulation after 1 hour of run time. Flow rates in five trials deviated from the mean by up to 10% (Fig 2A, black error bars marking two standard deviations). Moreover, the first chamber showed anomalously low flow rate compared to the target (Fig 2A). Consequently, the eight chambers showed large standard deviation (Fig 2A, blue error bar) when run as replicates.

**Fig. 2.**
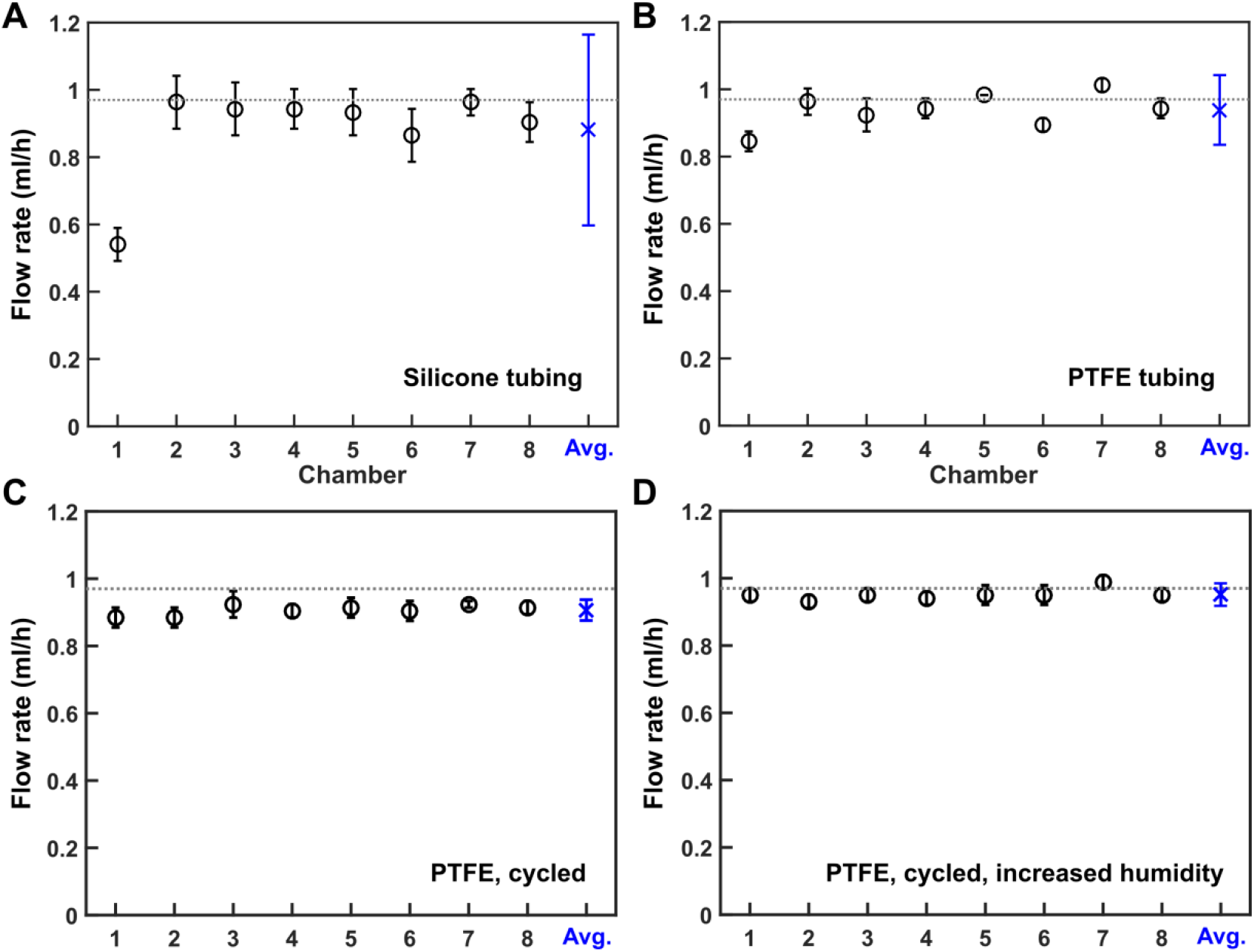
Chemostat flow rate improvements. Flow rates were measured by weighing media accumulated in chambers after 1 hour of run time, with positive pressure on and media outlets disabled (i.e. outlets elevated so that no medium flows out). Data point for each chamber represents 5 trials, with error bars indicating two standard deviations. The average and the two standard deviations of flow rates of all eight chambers are marked in blue. Target flow rate, 0.972 ml/h, is marked by a grey line. (**A**) In the original Klavins Lab design, the first chamber showed anomalously low flow rate. Within Chambers 2 - 8, the one-hour flow rate could vary from the mean by as much as 10%. (**B**) Replacing silicone tubing with PTFE tubing, (**C**) cycling dispense order, and (**D**) increasing the humidity of inflow air improved the consistency and accuracy of flow rate in terms of smaller error bars and closer match to the target. The average flow rate (blue) is 0.95 ml/h, with a two standard deviation of 0.03 ml/h.

### Rigid tubing reduces the flow rate anomaly of the first chamber

What might cause the anomalously slow flow rate in Chamber 1? We reasoned that this might result from the medium delivery silicone tubing expanding or shrinking as the internal pressure changes. The silicone tubing (green in Fig. 1B) is elastic ^7^ due to its relatively small Young’s modulus (typically between 0.005 GPa and 0.02 GPa). The elasticity allows pinch valves to work, but may also allow the tubing to change its inner diameter. This can affect flow rate, as we discuss below.

Specifically, at the beginning of each operation cycle, the pinch valve to reservoir is opened and the syringe pump draws fresh medium. The pressure at an arbitrary point *z* in the tubing is *P*(*z*) = *P_0_* − *ρgh_0_* (Fig 3A). Here, *P_0_* is the atmospheric pressure present in the reservoir, and *ρgh_0_* is the hydrostatic pressure of the media in the tubing, where *ρ* is the density of the media, *g* is Newton’s constant of acceleration at the surface of the earth, and *h_0_* is the vertical distance between the surface of the media in the reservoir and the location *z* (Fig 3A). When syringe pump delivers medium to Chamber 1, the pinch valve to Chamber 1 is opened. The pressure at point z becomes 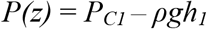 (Fig 3B). Here, *P_C1_* is the pressure in Chamber 1, and since positive pressure is applied, *P_C1_* > *P_0_*. Furthermore, *h_1_*, the vertical distance between the liquid surface of Chamber 1 medium inlet and *z*, is typically less than *h_0_* in our experiments. Thus, the pressure at point *z* can increase significantly when the syringe pump switches from the reservoir to Chamber 1 (Fig 3B, Δ*P*_Re*s*,−>*C*1_ > 0). As the liquid level in reservoir drops, *h_0_* increases, and the pressure increase from switching from Reservoir to Chamber 1 becomes even more drastic. Positive Δ*P*_Re*s*−>*C*1_ causes the tubing to expand, and results in the lower-than-expected medium input into Chamber 1 (Fig 2A). In contrast, when the pump switches from Chamber 1 to Chamber 2, pressure change (Fig 3C, Δ*P*_*C*1−>*C*2_) will be small due to the similarity in air pressure and inlet liquid height between the two chambers. Consequently, chambers subsequent to Chamber 1 do not display as large a deviation as Chamber 1 (Fig 2A).

**Fig. 3.**
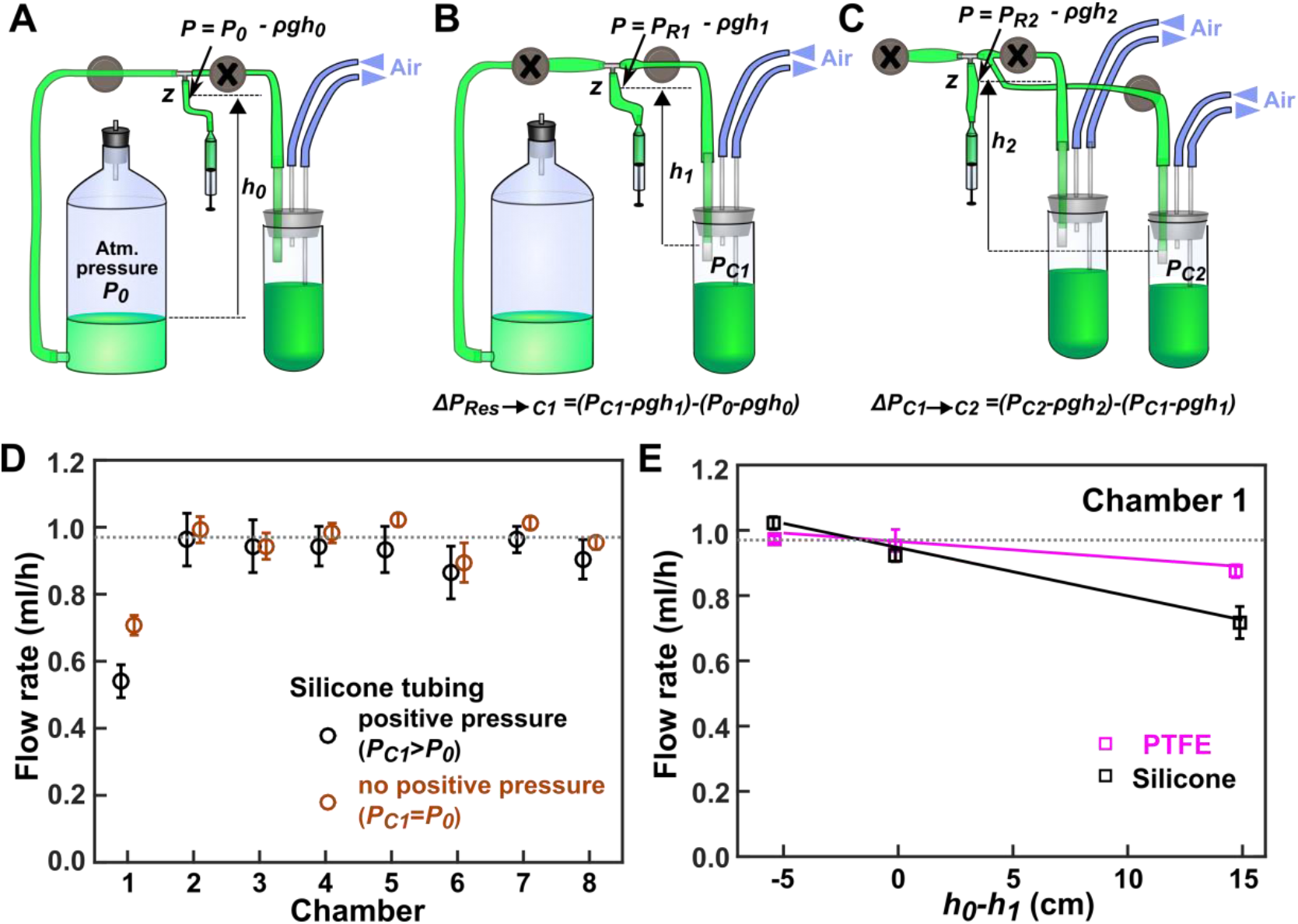
Pressure changes during medium delivery can lead to silicone tubing expansion and flow rate reduction. (**A**) For the syringe pump to draw medium from reservoir, we open the pinch valve to the reservoir (grey circle without cross) and leave pinch valves to eight chambers closed (here, pinch valve to the first chamber is shown; the rest not shown). The pressure at an arbitrary location z in the tubing is *P_Reservoir_(z)* = *P_0_* – *ρgh_0_*. See main text for explanation. (**B**) When the pinch valve to Chamber 1 is open (and the rest pinch valves having been closed), the pressure becomes *P_C1_* – *ρgh_1_*. Since positive pressure is applied to all chambers, *P_C1_* > *P_0_*. Under our typical setup, *h_0_*>*h_1_*, and as the reservoir media surface level drops during cultivation, *h_0_* will continuously increase. Thus, pressure in z increases as the pinch valve to Chamber 1 opens, causing tubing to expand. Consequently, a certain volume of medium that should have been dispensed to Chamber 1 instead remains in the stretched tubing. (**C**) Pressure changes between chambers is much smaller than that between the reservoir and Chamber 1. This is because positive pressure and liquid surface level are similar among chambers (e.g. *P_C1_* ~ *P_C2;_ h_1_* ~ *h_2_*). (**D**) Removing positive pressure from the chambers partially remedies the abnormally low flow rate of Chamber 1, and reduces the variability of the flow rate in each chamber. Flow rates were measured over 1 hour intervals. N=3. (**E**) Altering the height of the reservoir liquid level changes the flow rate of the first chamber. We adjusted the reservoir altitude to alter *h_0_*–*h_1_* and measured flow rate. When silicone tubing was used (black), we obtained a slope of -0.014+/-0.003 ml/h per cm height difference. This effect was reduced by a factor of 3 (magenta, slope= -0.0050+/-0.0001) after replacing silicone tubing with PTFE tubing (except for short sections where pinch valves require silicone tubing). In these measurements, positive pressure was absent. The data for both D and E were obtained utilizing 33 dispense cycles/h. Since each cycle introduces a fixed volume deficit, more cycles per hour means greater error in chamber 1.

If the above reasoning is correct, then reducing the pressure change Δ*P*_Re*s*−>*C*1_ by turning off air pumps (*i.e. P*_*C*1_ = *P*_0_) should mitigate the anomalous flow rate of Chamber 1 while only minimally affecting other chambers. This was indeed the case (Fig 3D, compare brown and black symbols). Furthermore, the flow rate became more consistent (Fig 3D, brown error bars smaller than black error bars), suggesting that unsteady air pressure contributed to variable flow rates when using silicone tubing.

Similarly, reservoir liquid level should affect flow rate of the first chamber. Indeed, when we adjusted the height of reservoir to achieve different levels of *h_0_-h_1_*, we observed a linear relationship between Chamber 1 flow rate and *h_0_-h_1_* (Fig 3E, black). When *h_0_-h_1_*=0, Chamber 1 flow rate was close to target (Fig 3E, black; Supplementary Fig 1). As expected, the flow rates of other chambers were not affected (Supplementary Fig 1).

To mitigate the tubing stretching problem, we replaced the entire tubing with PTFE tubing (2-mm inner diameter; 3mm outer diameter), except for 7cm-segments of the silicone tubing supplied by the valve manufacturer to be used with pinch valves. PTFE has a Young’s modulus between 0.40 GPa and 0.55 GPa ^8^, making it an order of magnitude more rigid than silicone. After tubing replacement, flow rate of the first chamber was much closer to the target across a range of *h_0_-h_1_* (Fig 2B, magenta). Furthermore, flow rate consistency improved in all chambers even with positive pressure (note smaller error bars in Fig 2B compared with Fig 2A).

Autoclaving our PTFE-modified device resulted in some leaks. These leaks occurred at barbed fittings inserted at various fluid branching points (Fig 1B, black “manifold”). To prevent leaks, a slice of silicone tubing gasket of ~1mm long was inserted between each barbed fitting and PTFE tubing (Supplementary Fig 2). This adjustment allowed at least five autoclaving cycles without any leaks, after which we formed new seal by replacing silicone tubing gasket and trimming the end of PTFE tubing.

### Modification of syringe pump operation for long-term culturing

We are currently using the NE500 pump (New Era Systems Inc.). Although the Klavins lab has a design for a 3-D printed pump, they also now recommend the NE500 because of its reliability.

With the original Klavins lab software, the syringe plunger would eventually collide with the end of syringe body (Fig 4A) after a few days of operation. We reasoned that a possible cause for collision was drift in plunger starting position over cycles. The pump would aspirate in one draw a sufficient amount of liquid for all eight chambers plus a backlash volume *b* (0.2 ml, see Supplementary Fig 3 for an explanation of backlash). Next, the pump would dispense the backlash volume *b* back to the reservoir, followed by dispenses into Chambers 1 through 8 (Fig 4B), in that order. Backlash is irrelevant for drift: as long as the threaded plunger drive rod (Supplementary Fig 3) turns by the same total amount in each direction there can be no drift. Rather, a single withdrawal volume versus multiple split dispense volumes may cause plunger to drift due to the necessity of converting volumes to integer numbers of steps (Fig 4B). Thus, we revised the original software so that we withdraw a series of volumes and then dispense them in the same order (Fig. 4C). This has indeed eliminated plunger drift and crash. Our solution has since been implemented by the Klavins lab.

Finally, we have modified the software so that if communication errors should occur between the software and the pump, the software will log errors and attempt to resend the command.

**Fig. 4.**
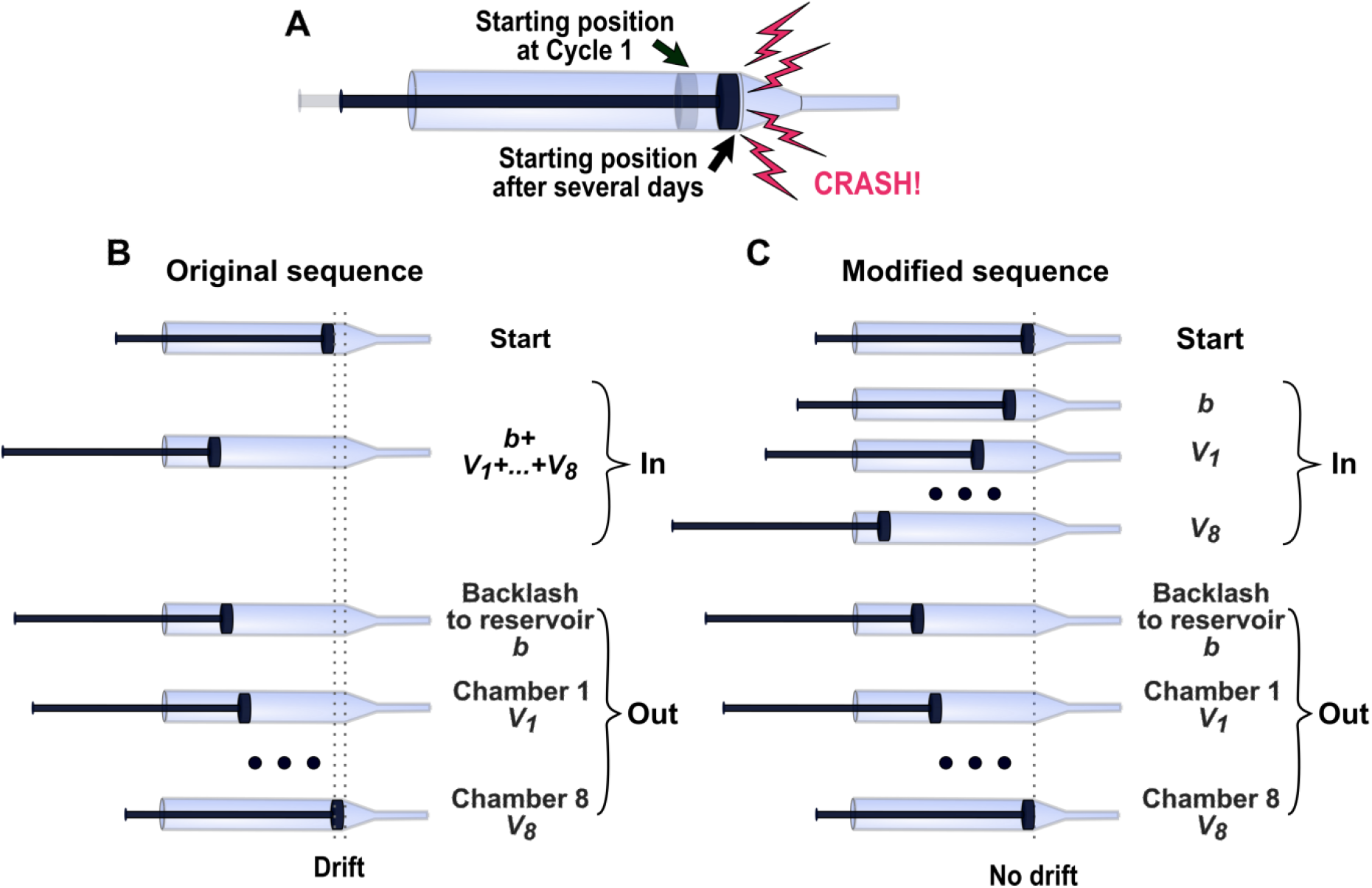
Identical liquid withdrawal and dispense sequences prevent syringe plunger from crashing into syringe body. (**A**) In the original system, we observed crashing of the syringe plunger with the end of syringe body due to slow drift in plunger starting position over time. (**B**) The Klavins lab software performed one withdrawal (In) and multiple dispenses (Out). Breaking up a single withdrawal volume into multiple dispense volumes may cause drift in plunger starting position. Here, *b* = volume to allow backlash to occur; *V_1_-V_8_*: volumes dispensed to Chambers 1 to 8. Volumes to different chambers can differ depending on flow rates desired. (**C**) In our modified software, a set of withdrawals are followed by an identical set of dispenses. This eliminates the drift in plunger starting point.

### Cycling dispense order increases flow rate consistency

Even after switching to PTFE tubing, statistically significant variations in flow rates were observed among chambers with the same target flow rate (Fig 2B). This could be caused by a variety of reasons including differences in positive pressures within different chambers or variable pump dispenses. We examined pump behavior by writing a LabView program to make the pump sequentially dispense a fixed volume. Indeed, dispensed volumes varied systematically in an order-dependent fashion (Fig 5). For example, the second and the fifth dispenses tended to under-deliver.

Given the systematic variations in dispense volumes (Fig 5) and the possibility of other order-dependent variations that affect flow rate (Fig 2B, Chamber 1), we modified the control software to cyclically permute the order of dispenses over the set of 8 chambers (Fig 6A). This should average out systematic variations in the syringe pump dispense volume. Moreover, any flow rate anomalies associated with switching from reservoir to chamber will be averaged over the eight chambers. This modification indeed reduced variations in flow rate among chambers whether positive pressure is applied (compare Fig 2B and Fig 2C) or not (compare Fig 6B and Fig 6C).

**Fig. 5.**
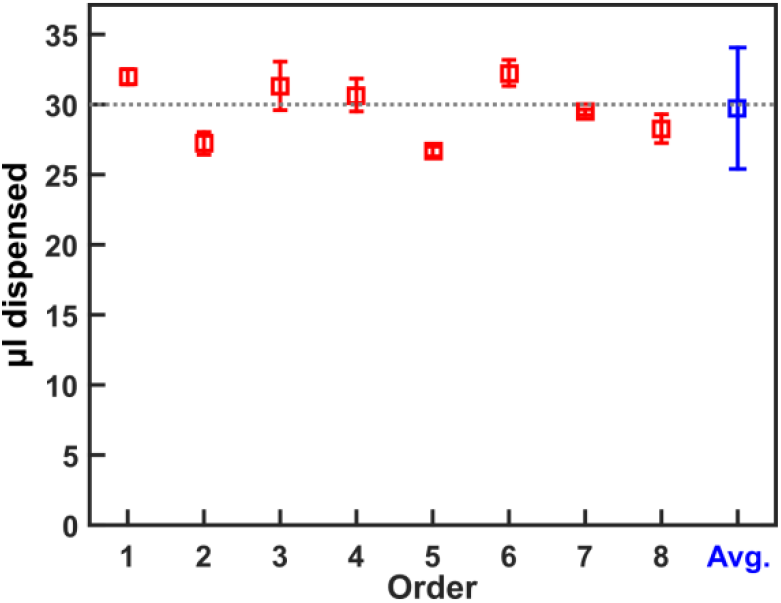
Order-dependent variations in syringe pump dispense amounts. We wrote a LabView program to test the pump. For each cycle, the pump withdrew H_2_O and made a sequence of eight 30μl-dispenses to a small vial. For each dispense, we weighed the vial before and after that dispense. Data from four cycles are plotted. The average dispense is 29.9 μl, closely matching the 30 μl target volume. However, order-dependent variations are clearly visible. For example, the second and the fifth dispenses are always below the target, while the first and sixth are always above. Error bars: 2 standard deviations.

**Fig. 6.**
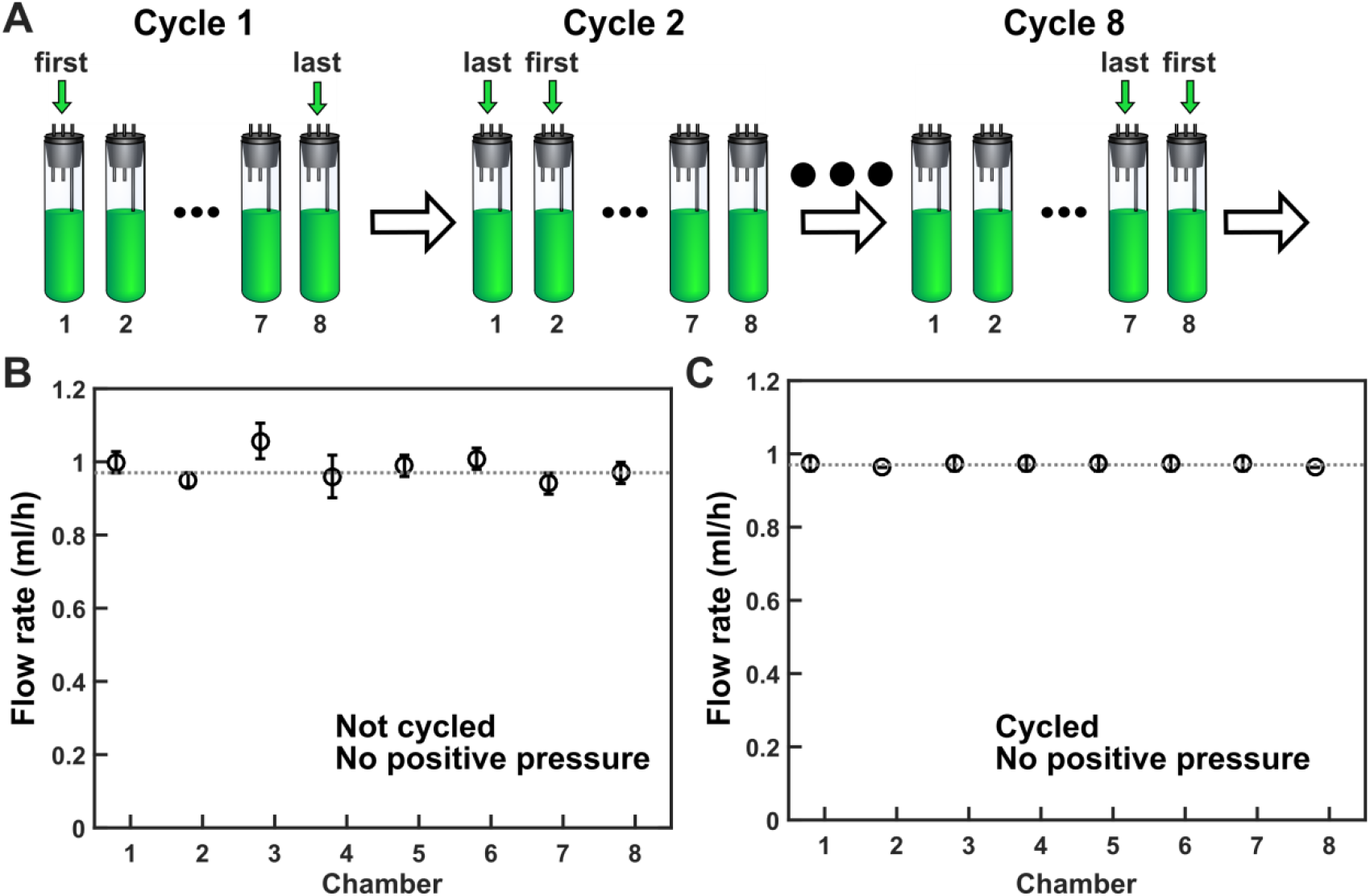
Cycling dispenses reduce variations in flow rate. (**A**) Cycling the sequence of dispenses over the eight chambers can average out variations every eight dispense cycles. (**B, C**) Cycling the sequence of dispenses indeed reduced flow rate variations among chambers. Here, we measured flow rates without applying positive pressure to reduce the effects of evaporation and the effects of different positive pressures within different chambers. N=5 and 3 for B and C, respectively.

### Increasing humidity of inflow air increases flow rate accuracy

After PTFE tubing replacement and cycling dispense order, we still observed a systematic deficit from the target flow rate (Fig. 2C). This is likely due to evaporation: a chamber with higher evaporation effectively receives a more concentrated medium at a lower flow rate. Consequently, cells will grow slower and reach a higher steady-state density (Equations 3 and 4).

To reduce evaporation, we have modified the original design which generates large air bubbles in a humidification vessel (Fig 7A). Air in those bubbles becomes humidified as water molecules evaporate from the bubble inner surface into the air inside the bubble. Air above the bubbler liquid surface is also humidified due to evaporation from the liquid surface. Because the air is continually flowing, it spends a limited amount of time in contact with the humidifying water, and may not become saturated with water vapor. Thus, we added an aquarium bubbler stone at the end of the submerged tubing to create a cloud of small bubbles (Fig 7B). We also maintained water depth in bubbler at a sufficient level. Increased surface area and sufficient resident time in water allow more water to evaporate into the inflow air. As a precaution we added aerosol trap to prevent tiny water droplets from entering and clogging air filters used to maintain the sterility of inflow air (Fig 7B). Humidifying inflow air indeed improved flow rate, bringing it closer to the target (Fig. 2D).

**Fig. 7.**
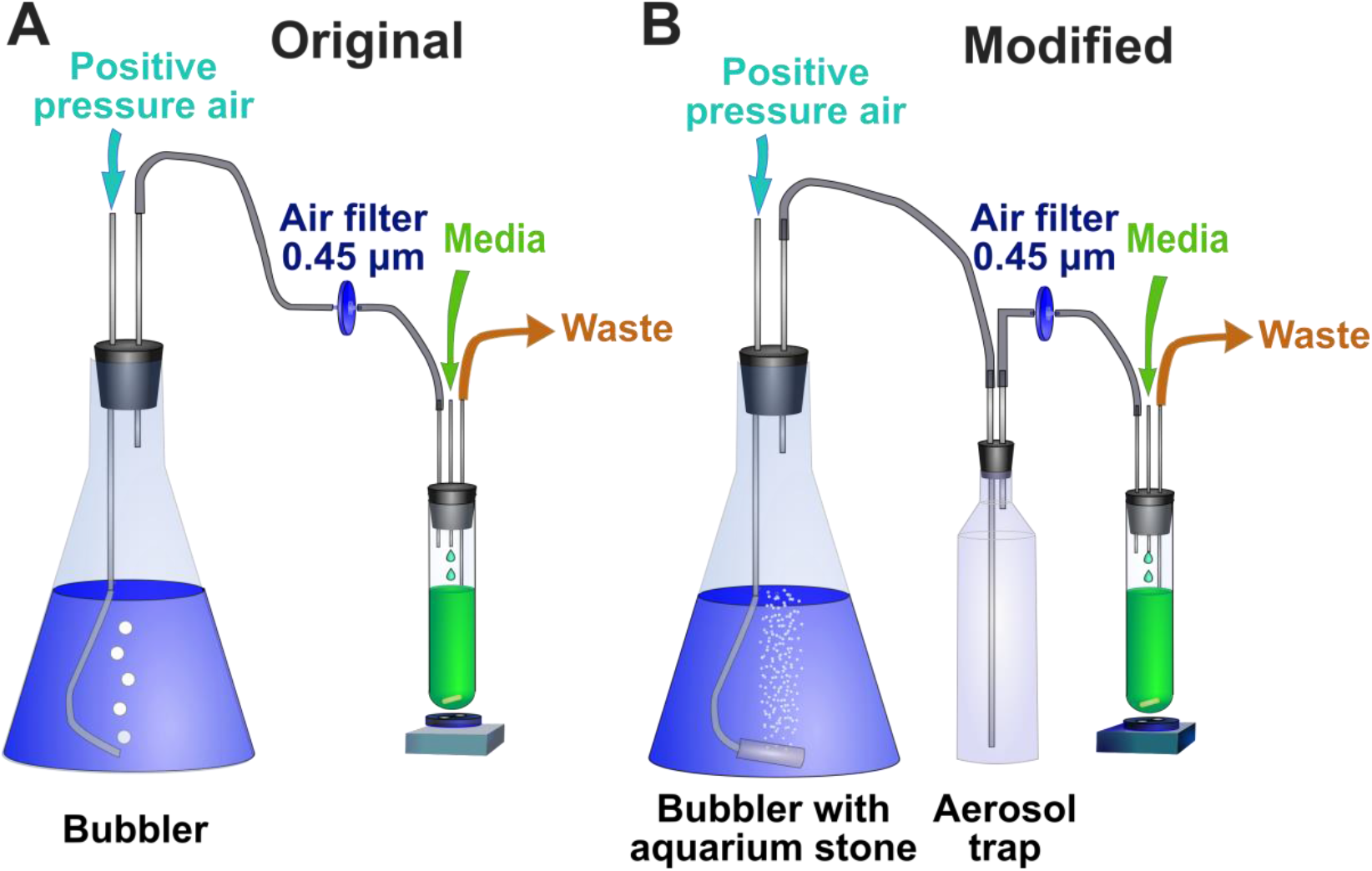
Increasing humidity in inflow air reduces evaporation. Humidified air flow creates positive pressure in chambers, forcing excess culture effluent out to waste. (**A**) In the original design, air humidification is limited due to large sizes of bubbles. (**B**) In our modified design, an aquarium stone generates numerous tiny bubbles. Liquid depth in bubbler is maintained at a sufficient level throughout the run so that air in bubbles has sufficient time to become humidified. The size of bubble cloud is indicative of airflow magnitude, and can be used to adjust airflow with a regulatory valve if needed.

Even with these modifications, actual flow rates are still lower than target values under positive pressure. Without any flow of liquid, the evaporation rate was measured to be 0.05 ml/h at 30°C, with a two standard deviation of 0.02 ml/h. With the flow of liquid, after carefully controlling for reservoir height and temperature, flow rates were 0.01 to 0.07 ml/h below targets in the tested range, and the value of deficit was largely independent of target value. For example, for 13h doubling time, the average deficit was 0.041 ml/h (2σ = 0.040 ml/h). For 3h doubling time, the deficit was 0.034 ml/h (2σ = 0.028 ml/h). The evaporation rate shows no significant dependence on flow rate. Thus, the percent deficit from target will be larger for slower flow rate.

### High consistency and accuracy under a range of target flow rates

We tested a range of flow rates in our chemostats. We found that our chemostats can reliably achieve 3h to 13h doubling times (Fig 8; Fig 2D). A shorter doubling time can in principle be achieved by using a larger-diameter syringe. A longer doubling time will be limited by the effects of evaporation and discontinuous flow. At a doubling time of 13h, about one drop of ~30μl is dispensed per ~111s. At a slower dilution rate, longer times will be required for each medium droplet to drip, which may affect cell physiology and increase flow rate variability due to evaporative differences between chambers. At extreme cases (e.g. doubling time of over a week), evaporation rate may equal to flow rate, and the chemostats will not be operational. Nevertheless, the operational ranges we have tested are sufficient to investigate over a four-fold difference in population growth rates, allowing researchers to examine physiological and evolutionary differences between mildly-starved versus strongly-starved *S. cerevisiae*, for example.

**Fig. 8.**
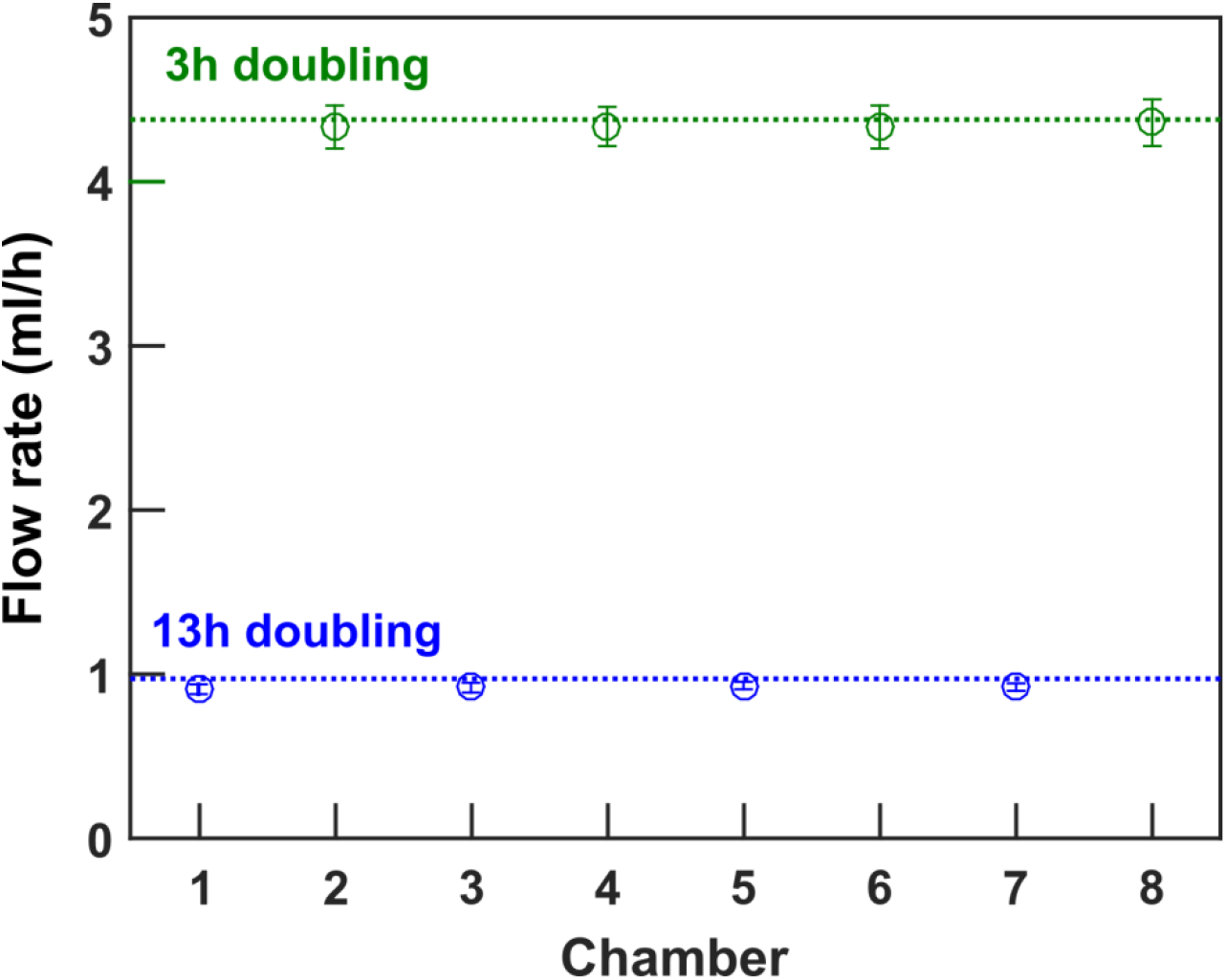
Accurate flow rates across a wide range. We ran four chemostats at 3h doubling time (green) and four chemostats at 13h doubling time (blue). The average flow rate is within 3% of target for 13 hour doubling time, and within 1% of target for 3 hour doubling time.

### Precise control of flow rate is important for phenotype quantification

Variations in flow (and thus growth) rate can profoundly affect experimental measurements. For example, Varma et al^9^ found that in a glucose-limited chemostat, *E. coli* released acetate as a byproduct only when dilution rate exceeded a certain threshold.

To test the importance of flow rate control, we cultured a purine-requiring, lysine-overproducing *S. cerevisiae* strain^10^ in hypoxanthine-limited chemostats run at a doubling time of 6h (dilution rate 0.116/h) or 7h (dilution rate 0.099/h). These corresponded to a dilution rate difference of ~16%, smaller than the deviation seen for Chamber 1 in the original design (Fig 2A). The dead:live cell ratio (and thus death rate; Equation 5 in Methods) was higher in 7h chemostats (severer nutrient limitation) than in 6h chemostats (Fig 9 A and B). Since dilution rate was lower in 7h chemostats, lysine accumulated to a higher level compared to 6h chemostats (Fig 9 C and D; Equation 6 in Methods), thus creating different extracellular environments. Without a precise flow rate, variations in dilution rate will interfere with experimental measurements by introducing large errors (Fig 9, B and D black). Thus, precise control of flow rate is critical for quantifying phenotypes (Fig 9 A and B) and the environment cells create (Fig 9 C and D).

**Fig. 9.**
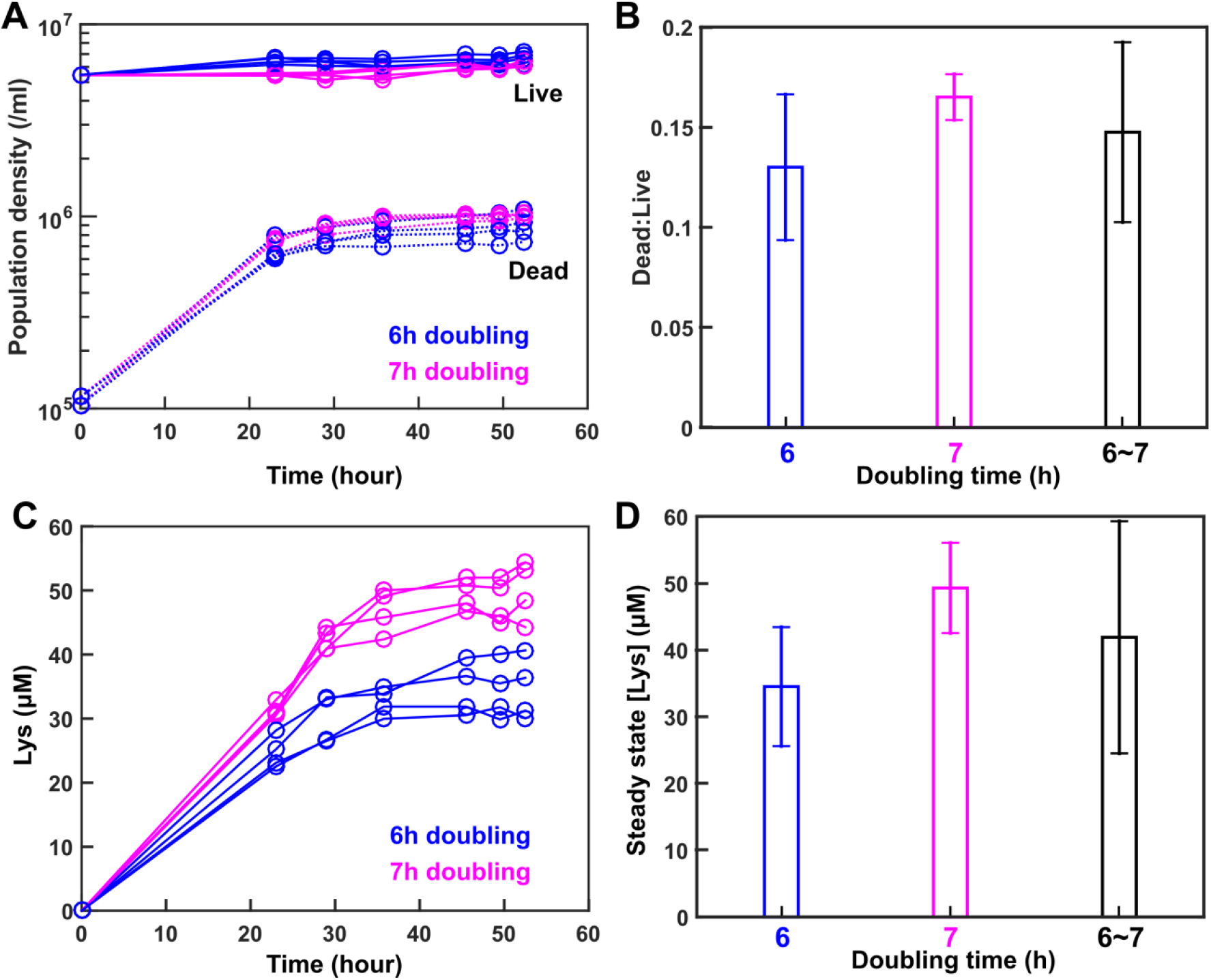
Precise flow rate is important. We ran four chemostats at 6h doubling time (blue) and four chemostats at 7h doubling time (magenta). (**A**) Dynamics of live and dead cell densities (Methods) reach steady states. (**B**) As expected, the ratio of dead cell density averaged over the last four time points to live cell density averaged over the same period of time (and thus death rate) was higher in 7h chemostats than in 6h chemostats (two-sample *t* test, P value = 0.01). (**C**) Dynamics of released lysine (Methods) reach steady states. (**D**) Steady-state lysine concentration (averaged over the last three time points) was higher in 7h chemostat compared to 6h chemostat (two-sample *t* test, P value = 0.002). In (**C**) and (**D**), black bar combines data from all eight chemostats. Error bars indicate two standard deviations.

## Summary

We have improved multiplexed turbidostats to run as chemostats without expensive feedback systems. Three major modifications are: (i) replacing silicone tubing with PTFE tubing to increase tubing rigidity (Fig 2B), (ii) permuting dispense order to overcome systematic errors of syringe pump dispenses (Fig 2C), and (iii) adding a bubbler stone to reduce evaporation (Fig 2D). After implementing these modifications, the flow rates of all eight chemostats, which can be independently adjusted, consistently match their respective target flow rates (Fig 2D; Fig 8). Our system can be used to quantify cell physiology and to carry out evolution experiments under nutrient limitation.

---

**Supplementary Fig 1.**
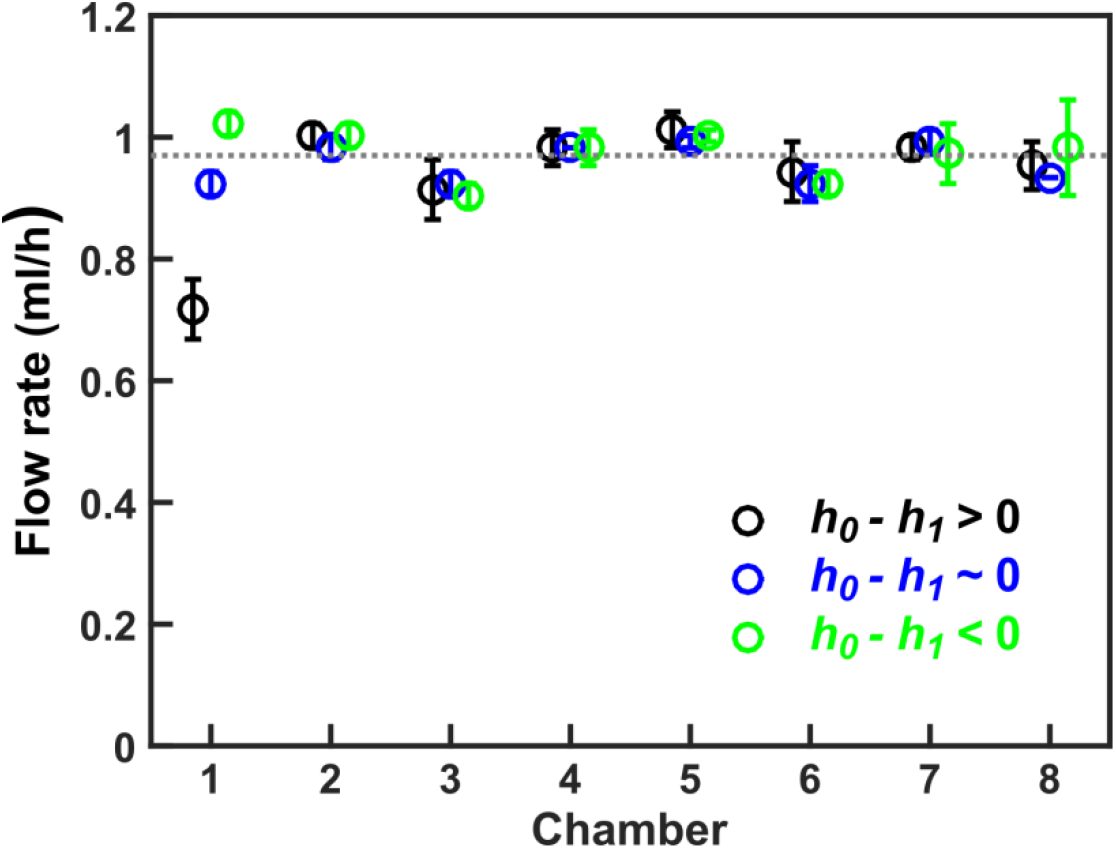
Variable flow rate in first chamber when using silicone tubing. Depending on the liquid level of reservoir (*h_0_*, which changes during the course of an experiment) and the liquid level of media input to Chamber 1 (*h_1_*), the first dispense (which is to Chamber 1) might deviate from the target (grey line). Chambers 2 to 8 are much less affected.

**Supplementary Fig 2.**
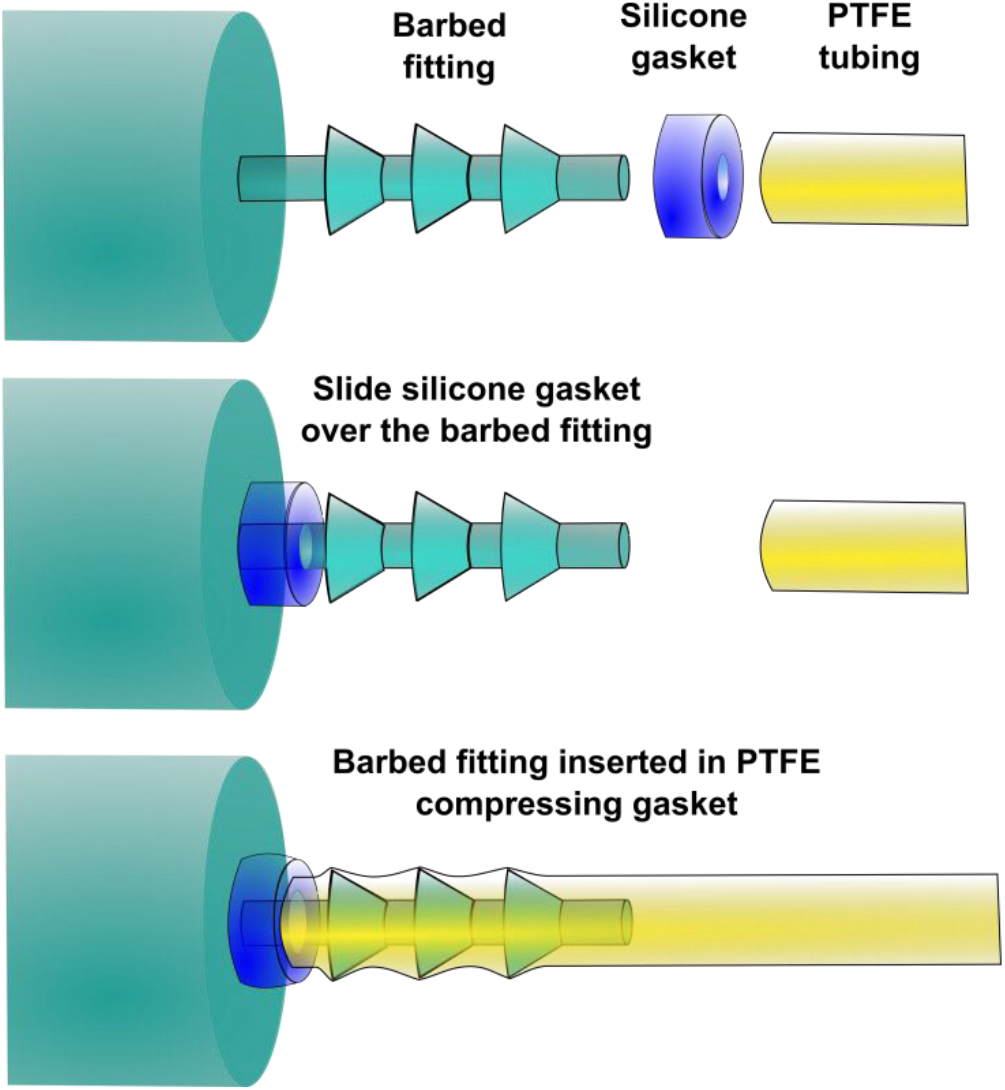
Silicone gasket assembly. To prevent leaks from the junction of rigid PTFE tubing with the barbed fittings of manifolds, we added a small silicone gasket. Because we could not find commercial sources for such small silicone gaskets, we cut approximately 1mm sections of silicone tubing (1/16” ID 1/8” OD from VWR International) with a single edged razor blade. The silicone gasket is placed over the barbed fitting, and the fitting is inserted into the tubing, pressing the end of the tubing against the gasket, thus forming a flexible seal. Silicone gasket assembly allows at least five cycles of autoclaving without leakage.

**Supplementary Fig 3.**
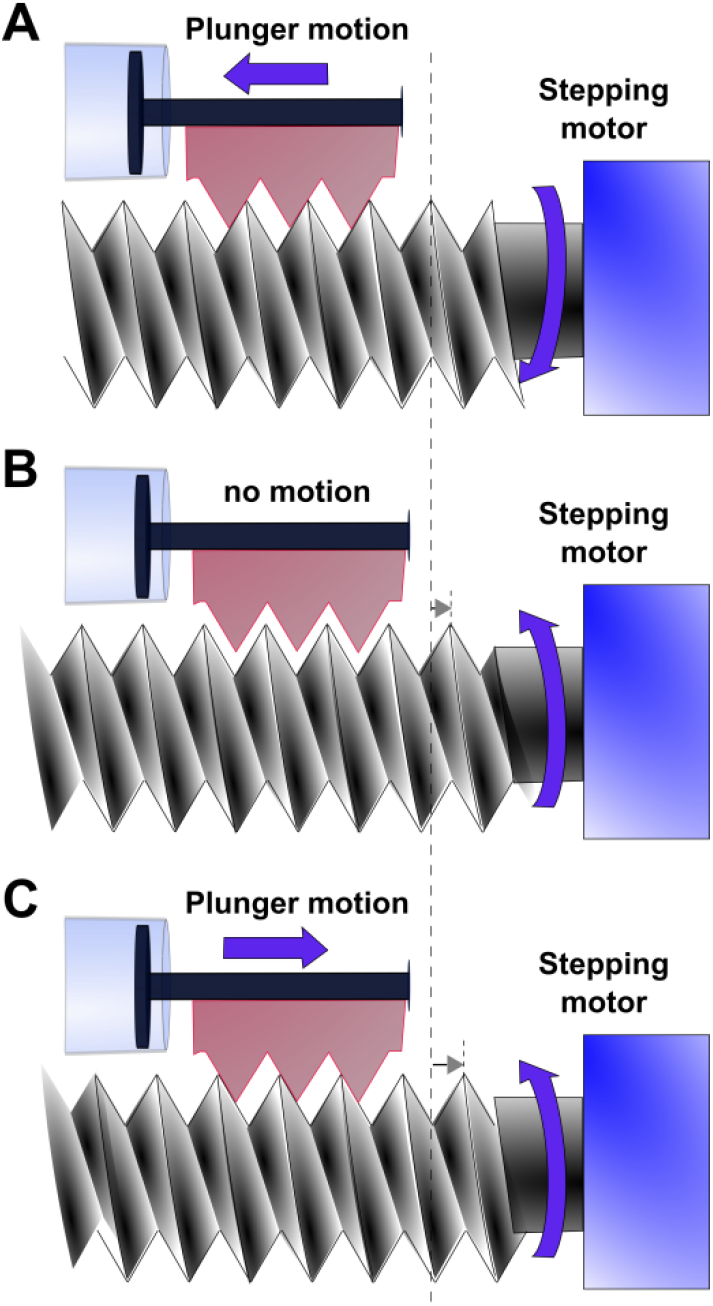
Pump backlash. The syringe pump uses a stepping motor and a threaded drive rod to move the syringe plunger. Backlash is defined here as the maximum turning of stepping motor which does not lead to plunger movement. (**A**) Counter-clockwise turning of stepping motor (viewed from the right-hand side) drives plunger to move left. (**B**) When the direction of turning is reversed, the threads of the plunger holder are detached from the threads of the motor drive screw. Thus, even as the motor turns, the plunger does not move. This is the backlash. (**C**) Re-engagement between plunger and stepping motor leads to plunger motion. Long dashed line marks a reference position, and short dotted lines mark one tooth of the stepping motor. ~0.2 ml was used for backlash correction volume.

## Materials and Methods

Instructions for construction of the Kalvins lab turbidostat, unrelated to our modifications, can be found at the Klavins lab website.

### Modified components

#### Syringes

A smaller syringe diameter will result in smaller variations in flow rate due to a longer travel distance of the plunger. However, a smaller syringe diameter requires a faster speed to deliver the same volume in the same amount of time. BD 1ml syringes (BD309659) have an inner diameter of 4.73 mm. The maximum pump speed that the NE500 will accept for this diameter is 53 ml/h. This flow rate could barely achieve a 4 hour doubling time in eight chambers with 19 ml volume, if the pump were to work continuously without latency. To see this, since withdrawal and dispense account for two operations, the maximal pump rate translates to 53 ml/h/(8 chambers*(19x2) ml/chamber) amounts to 0.174/h, which is about 3.98h doubling time. Pump latency (e.g. a backlash compensation volume per cycle; waiting time for the pinch valves to open and close; delays between syringe pump operations to allow pressure equalization) means that a doubling time of even 8h cannot be achieved in this system with a 1ml B-D syringe. Because of the improvements we have made, we could use a 3ml EXELINT^®^ syringe (Ref. 26102) to achieve a doubling time of 3~13 h with high precision. At a slower flow rate, each cycle involves a smaller displacement of the syringe plunger, and thus pump variations become more prominent. We measured the error of flow rate at 13h doubling time (N=5). In the absence of positive pressure (Fig 6B), two coefficient of variation among 8 chambers is 1%, and the maximum deviation from the target flow rate is -2%. With positive pressure (Fig 2D), two coefficient of variation among 8 chambers is 3%, and the average and the maximum deviation from the target flow rate is -2% and -4%, respectively.

#### PTFE tubing

The PTFE tubing used in our setup was generic tubing purchased on eBay, with no manufacturer tolerance specifications. Measurements with a digital caliper gave an outer diameter of 3.07 mm, with a two standard deviation of 0.06 mm, and a wall thickness of 0.59 mm with a two standard deviation of 0.17 mm. Measurements were performed on 5 separated sections of tubing, at 8 different angles to average out ellipticity and other irregularities. To connect PTFE with manifold barbed fitting, we use a silicone gasket fitting (Supplementary Figure 2).

#### Aquarium bubbler stone and regulation valves

We used 1” Aircore Sand Airstone (JW Pet Company, available on Amazon) in the humidification vessel. Any aquarium bubbler stone should give similar results. We used Airline Control Kit (Lee’s Aquarium & Pet Products, available on Amazon) as air flow regulator.

### Measurements of flow rates and distances

Flow rates in the eight chambers were measured by running the chemostat software for 33 cycles of 111 seconds per cycle, or 1.02h. The waste tube was well above the liquid level in the chamber to prevent liquid outflow. This allowed us to measure flow rate by quantifying the rate of increase in chamber weight, using a Mettler PE 3600 balance. Each chamber was initially filled with a volume approximating the typical working volume of 19 ml, weighed, and capped with a temporary rubber stopper (without tubing) to limit evaporation. Bubbles were purged from chemostat lines by running water to a waste container (all chemostat stoppers were open to the air). Afterwards, the temporary stoppers were switched to chemostat chamber stoppers, and chambers were placed in their holders. A timer was set to 1h to alert the operator to the initiation of the 33^rd^ cycle so that chemostat software could be stopped at the end of that cycle. The chambers were then removed, and capped with temporary stoppers. Each chamber was weighed without stopper, and the difference in weight was used to calculate the volume of water pumped into the chamber (assuming the density of water = 1.00g/cm^3^ at 30°C). The flow rate was determined by dividing the calculated volume by the 1.02h total operation time. Tests with positive pressure (Fig 2) were performed with the air pumps on. Otherwise, the air pumps were off.

Distances were measured using a digital caliper (Fisher Scientific 8 inch). Sighting across caliper jaws is required for measuring liquid height within each chamber. Multiple measurements were averaged.

### Operation of chemostat

#### Chemostat set up

We assembled the chemostat tubing, chambers, and empty media reservoir, and covered open tubing endings with Leuer locks or foil caps. We then autoclaved the entire assembly, skipping the vacuum dry cycle to preserve the integrity of tubing attachments. After autoclaving, we carefully uncovered these open ports and attached them to air filters (Fig 7B) or the syringe pump (Fig 1B). We autoclaved air filters separately with a vacuum cycle to ensure that they stayed dry and fully functioning, and fit the air filters over the bubbler ports to ensure sterility of inflow air. We then sterile filtered a sufficient amount of SD + 20 μM hypoxanthine into the reservoir for the desired experiment length. For example at 6h doubling time, since each chamber is of 19 ml, the flow rate will be 19*ln(2)/6= 2.195 ml/h per chamber. Thus, if all eight chambers were used over 60 hours, then we would need at least 2.195 ml/h/chamber*8 chambers *60 h = 1053 ml. When medium refill is necessary, the program could be momentarily stopped to switch to a new reservoir of media. Alternatively, we have used sterile tubing to drain fresh media into the reservoir without stopping the software.

We have set pump dispense cycle time to be 111 sec. Thus, each chamber gets about 2.195 ml/(3600 sec)*111 sec= 67.7 μl per cycle. For each cycle, we need to retrieve from reservoir 67.7 μl /chamber*8 chambers = 0.541 ml. We take out an additional ~0.2ml to correct for backlash (Supplementary Figure 3).

We allowed the medium to flow through and fill the tubing and syringe but not the chambers by sending commands to individually open pinch valves to allow the media to fill the tubing, but closing the valve when medium starts to drip into the chamber. If a bubble should form at near the barbed fitting, pressing the tubing against the fitting can reseal the junction. Throughout the run, we ensure that liquid level in bubbler maintains a depth of at least 7.5cm, so that the bubbles have sufficient time to humidify.

#### Chemostat inoculation

We used strain WY1340 which is in the RM11 background, expresses GFP, requires adenine or hypoxanthine, and overproduces and releases lysine (*ho::loxP AMN1-BY ste3::NAT fba1::FBA1-EGFP-loxP ade8::loxP lys21::LYS21^o/e^*). We steaked out frozen stock on rich medium YPD (10 g/L yeast extract, 20 g/L peptone, 20 g/L glucose) with 100 μM supplemental hypoxanthine. After allowing two days for colonies to grow up at 30°C, we inoculated an isolated colony in defined minimal medium SD (6.7 g/L Difco^TM^ yeast nitrogen base w/o amino acids, 20 g/L glucose) with 100 μM supplemental hypoxanthine (non-limiting). We allowed this culture to grow overnight at 30°C and harvested exponential phase cells (7x10^6^ ~3x10^7^/ml). We washed away supplemental hypoxanthine by spinning down cells and re-suspending them in SD. We then pre-starved the cells for 24h at 30°C to deplete hypoxanthine storage, taking care to dilute to <7x10^6^/ml so that cells would not be limited for any other resource besides hypoxanthine while undergoing up to 5-fold residual growth.

After cells had starved for 24h, we measured cell density using a flow cytometer and diluted cells with SD to roughly the expected steady state density. For example, 20 μM hypoxanthine in the input medium yields a steady-state cell density of 5~7x10^6^ cells/ml. Using a sterile 30 ml syringe, we filled each chemostat with the diluted cell culture. We programmed the chemostat to run four chambers at a 6h doubling time (~68 μl dripped in every 111s) and four chambers at a 7h doubling time (~58 μl dripped in every 111s).

To allow cells to reach a steady-state density and physiology, we allowed chemostats to run for 23 hours before sampling. At each time point, we withdrew ~1.5ml of culture using a sterile syringe. We immediately filtered 1ml through a 0.45 μm filter and froze the supernatant at −80°C for later lysine analysis. We used the remaining 0.5ml to measure cell densities using flow cytometry.

#### Quantifying chemostat dynamics

In order to measure live and dead cell densities in each sample, we mixed it with Fluoro-Max^TM^ red fluorescent beads (Thermo Fisher Cat. #R0300) of a known concentration, and a 12,500x dilution of 1mM To-Pro3 (nucleic acid stain for dead cells). We ran this mixture on a flow cytometer. Because WY1340 expresses GFP and because dead cells are stained by To-Pro3, we identified GFP cells using 50 mW 488nm laser excitation with 505 ± 5 nm filter (BluFL1) and ToPro+ dead cells using 25 mW 637 nm laser excitation with 660 ± 8 nm filter (RedFL1). Beads were identified as high in both 50 mW 407 nm laser excitation with 450 ± 25 nm filter (VioFL1) and 50 mW 488nm laser excitation with 530 ± 15 nm filter (BluFL2). We then used cell:bead ratio and bead stock concentration to calculate live and dead cell density.

To measure lysine concentration, we employed a bioassay using a lysine-requiring yeast strain ^10^. We mixed 120 μl of each unknown sample with 30 μl of a master mix containing lysine-requiring cells at a low density (<1x10^5^/ml) and 5x SD to ensure that no other metabolites are limiting. Within each assay, we also used SD medium supplemented with various known concentrations of lysine to establish a standard curve. We placed all samples in a 96-well plate which we wrapped with parafilm, and allowed cells to grow to saturation at 30°C for ≥24h. We re-suspended cells using a Thermo Scientific Teleshake (setting #5 for ~1 min) and read culture turbidity using a BioTek Synergy MX plate reader. Since final turbidity of the lysine-requiring strain correlated linearly with lysine concentration up to a point in the standard curve, we could use the standard curve to infer metabolite concentrations of unknown samples.

#### Death rates and metabolite concentrations in chemostats

Chemostat dynamics can be described as

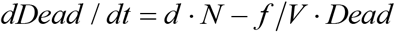

where *Dead* is the dead population density, *N* is the live population density, and *d* and *f/V* are death and dilution rates, respectively. At steady state,

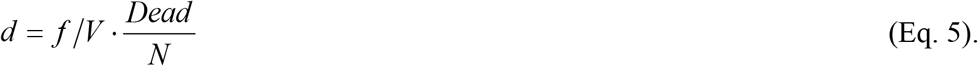

That is, death rate is the dilution rate multiplied by the ratio of dead to live population densities. If live cells release lysine at a constant rate *re*, then

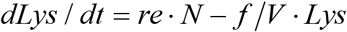

where *Lys* is the lysine concentration in culturing vessel. At steady state,

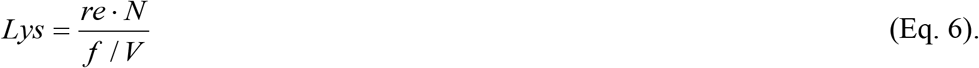

Thus, a higher dilution rate *f/V* leads to a lower concentration of released lysine.

## Acknowledgements

We thank Jean-Paul Toussaint and Chris Takahashi for their help in the initial fabrication and assembly of the turbidostat array, and Kennan Mell for testing and help with software implementation. We thank Chris Takahashi and current members of the Shou Lab (Robin Green, Li Xie, and Alex Yuan) for productive discussions of this project. This work is funded by the NIH, the W.M. Keck foundation, and Fred Hutch Cancer Research Center.

